# Optimized analysis for sensitive detection and analysis of single proteins via interferometric scattering microscopy

**DOI:** 10.1101/2021.08.16.456463

**Authors:** Houman Mirzaalian Dastjerdi, Mahyar Dahmardeh, André Gemeinhardt, Reza Gholami Mahmoodabadi, Harald Köstler, Vahid Sandoghdar

## Abstract

It has been shown that interferometric detection of Rayleigh scattering (iSCAT) can reach an exquisite sensitivity for label-free detection of nano-matter, down to single proteins. The sensitivity of iSCAT detection is intrinsically limited by shot noise, which can be indefinitely improved by employing higher illumination power or longer integration times. In practice, however, a large speckle-like background and technical issues in the experimental setup limit the attainable signal-to-noise ratio. Strategies and algorithms in data analysis are, thus, crucial for extracting quantitative results from weak signals, e.g. regarding the mass (size) of the detected nano-objects or their positions. In this article, we elaborate on some algorithms for processing iSCAT data and identify some key technical as well as conceptual issues that have to be considered when recording and interpreting the data. The discussed methods and analyses are made available in the extensive python-based platform, PiSCAT^§^.

---

Detection, characterization and imaging of small amounts of matter is of central importance in a large number of scientific and technological applications with biochemical and atmospheric analytics representing just two important examples (1). While techniques such as electron microscopy (2) or mass spectrometry (3) allow ultrasensitive detection of matter, the recent emergence of nanoscience and nanotechnology has rekindled the interest in achieving the same goals via optical techniques, leading to many breakthroughs such as scanning near-field optical microscopy, single-molecule spectroscopy, and super-resolution microscopy (4).

Optical detection of small nano-objects is often achieved via fluorescence, but the problems associated with labeling and photobleaching limit the scope of this approach. A powerful alternative is presented by the ubiquitous process of Rayleigh scattering, which is commonly detected via dark-field microscopy. The application of this method, however, is usually limited to the detection of particles that are larger than a few tens of nanometers. In 2004, interferometric scattering (iSCAT) microscopy was introduced (5; 6), which has since reached a remarkable real-time detection sensitivity down to single unlabeled proteins (7; 8; 9; 10). In our study, we present an optimized analysis platform with an emphasis on the prospects of pushing the limits of mass sensitivity further.

## 1. Fundamentals of iSCAT imaging

In iSCAT, one embraces the illumination field instead of avoiding it, as is done in darkfield microscopy. The illumination field, or a fraction of it, acts as a reference field that is interfered with the Rayleigh-scattered light of the sample (6; 11) such that the intensity on the detector reads

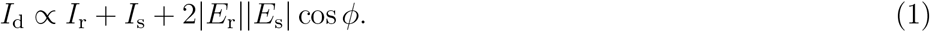

Here *E*_r_, *I*_r_ = |*E*_r_|^2^, *E*_s_, and *I*_s_ = |*E*_s_|^2^ denote the electric fields and intensities of the reference and the scattered light on the detector, respectively. The phase *ϕ* stands for the relative phase between the two fields, which usually has two main components: a Gouy phase due to the tight focus of an imaging arrangement and a traveling phase component stemming from the axial position of the particle. As shown in figure 1(a), the most common configuration of iSCAT operates in the wide-field reflection mode, where the light reflected from the sample interface acts as the reference field *E*_r_, and *E*_s_ is collected via the same microscope objective that delivers the illumination. However, iSCAT imaging can also be achieved in several other illumination and collection arrangements (6).

**Figure 1.**
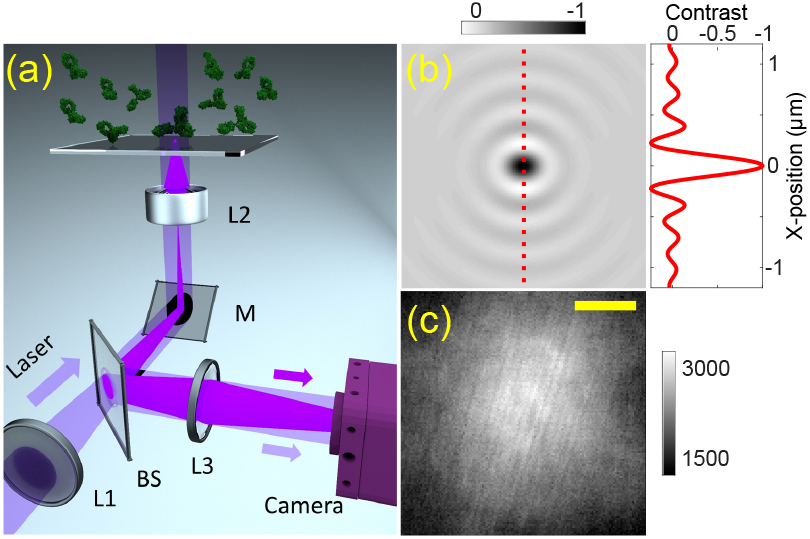
a) Schematics of the optical setup used in this work. A laser beam at a wavelength of 445 nm is focused onto the back focal plane of a high-NA objective resulting in a collimated beam at the sample position (diameter ca. 6 *μ*m). The beam is incident on a sample chamber consisting of a microscope coverglass sealed to the bottom of a plexiglass dish containing about 1ml of buffer solution. *L*_1_, *L*_2_, and *L*_3_ denote the wide-field, objective and imaging lenses, respectively. BS and M signify a beam splitter and a coupling mirror, respectively. b) Example of a synthetic iSCAT point-spread function (iPSF). The red curve presents the cross section marked by the dotted line. c) A raw frame of the background speckle without any particles in the field of view. Scale bar in (c) correspond to 1 *μ*m.

The iSCAT point-spread function (iPSF) of a wide-field setup results from the interference of a quasi-spherical wave emitted by the nanoparticle under study and a quasi-plane wave of the reference field (12). Figure 1(b) depicts a computed iPSF for a dielectric nanoparticle placed at the glass-water interface and in the focus of the microscope objective. A cross section (see red curve) shows that an iPSF in a wide-field setup contains several interference rings, the details of which depend on the focusing parameters and the exact particle position along the optical axis. In the presence of a limited signal-to-noise ratio (SNR), however, the higher-order rings are often not pronounced. Indeed, the large majority of iSCAT works have analyzed the central iPSF lobe (5; 7) although the rich ring structure of the iPSF has recently been exploited for 3D particle tracking (12; 13).

The key added value of iSCAT microscopy over conventional interferometric modalities is the detection of subwavelength nanoparticles (6; 11). The scattering cross section of such a particle illuminated at vacuum wavelength λ is given by (6)

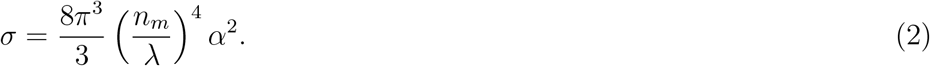

Here *α* denotes the polarizability of the nanoparticle given by

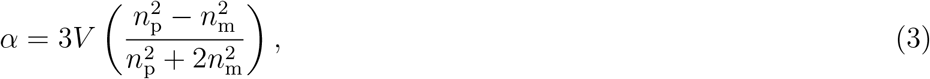

where *n*_p_ and *V* denote the refractive index and the volume of the particle surrounded by a medium of refractive index *n*_m_. Noting that 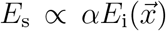 for an incident field 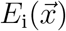 at particle position 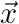, it becomes clear that the second term in equation (1) scales as *V*^2^ whereas the third (extinction) term scales proportionally with *V*. Thus, *I*_s_ in equation (1) becomes negligible for smaller particles so that the contrast (*C*) on the camera can be formulated as,

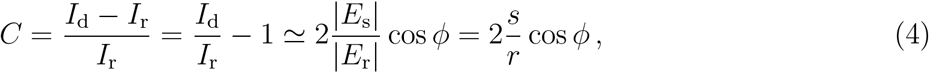

where *s* and *r* denote the proportionality factors between *E*_s_/*E*_i_ and *E*_r_/*E*_i_, respectively. Thus, *C* can be manipulated by the strength of the reference field. However, it should be kept in mind that the decisive metric for the detection sensitivity is the signal-to-noise ratio (SNR) and not *C* (5; 6; 11). In what follows, we consider the various components of the signal analysis for a typical sensing scenario, in which proteins are delivered to a chamber and precipitate on the bottom substrate positioned at the focus of an iSCAT microscope (see figure 1(a)).

## 2. Signal analysis

In the absence of extrinsic noise or signal fluctuations, one could detect arbitrarily small contrasts, only limited by the shot noise. Thus, considering that scattering is a linear non-saturable process, the SNR can be enhanced if one increases the incident power or the integration time. Scientific CMOS devices with quantum efficiencies up to 90%, fill factors close to 100% (back-illuminated CMOS chips), and pixel well depths ranging from tens of thousands to millions of electrons are well suited for iSCAT imaging although currently no camera possesses all of these features at the same time. For a pixel with a full well depth of 2 x 10^5^ electrons and readout noise of 220 e, the total shot noise in a single camera frame amounts to 2.2 x 10^−3^ of the signal level at saturation. This noise level is comparable to the iSCAT contrast of a 400 kDa protein placed at the glass-water interface (see SI. 1.1).

The first iSCAT sensing experiment with single-protein sensitivity reported a limit of about 66 kDa, corresponding to an iSCAT contrast of 3 x 10^−4^ in the arrangement shown in figure 1(a) (7). To reach this detection limit, one has to sum the signal of many frames. Although this is a straightforward conceptual procedure, its realization confronts several technical issues and a number of noise sources, stemming from the detector (readout noise, dark current, fixed pattern noise), laser intensity fluctuations (instrumental power fluctuations and shot noise), and phase noise caused by drifts and vibrations of the sample stage or the optical components, as well as the motion of the analyte particle. In other words, in addition to the shot noise of the optical field, iSCAT imaging faces several other sources of signal fluctuations that have to be addressed.

### 2.1. Power normalization

Available light sources suffer from intensity fluctuations, which are often larger than 0. 1%. In experiments on the detection of single dye molecules via extinction, these power fluctuations were corrected for by using a balanced photodetector (14). Here, a fraction of the incident light was measured on a second detector, acting as a reference for power normalization (PN). In a wide-field experiment, the sum of the signal over all the pixels in the field of view (FOV) provides a convenient and reliable intrinsic reference for PN (7).

### 2.2. Digitization noise

The next technical feature to be considered is the bit depth of the camera’s analog-to-digital converter (ADC). An *N*-bit ADC digitizes an analogue signal with the maximum amplitude into 2^*N*^ discrete levels. Thus, a commonly used monochrome 12-bit pixel has 4096 discretized gray levels, so that the smallest resolvable change in each frame corresponds to 2^−*N*^ ≃ 2.4 x 10^−4^. While this is much smaller than the shot noise of the full well, it is comparable to the contrast of a small protein. Thus, one might wonder if the digitization noise would impose a harsh limit on the detection sensitivity. However, because the noise in the signal projects it to the lower or higher neighboring values, averaging a large number of frames will converge the output to a higher resolution(15).

### 2.3. Differential treatment of images to eliminate image inhomogeneities

The most fundamental source of fluctuations in iSCAT microscopy stems from the signal of material features, such as surface roughness, adsorbates or other nanoparticles that generates a speckle-like background in the FOV. This was indeed the case in the early iSCAT measurements, where spin-coated stationary gold nanoparticles of diameter 5 nm were hard to distinguish from the background (5; 16). In the case of moving particles, the background can be eliminated by a differential treatment of the consecutive images of a video stream (7; 9; 17). This analysis can be done at the level of individual frames or batches of averaged frames, depending on the speed of the events of interest and the SNR in each frame.

Figure 1(c) shows an example of the raw camera image before proteins are introduced. The large bright spot indicates the extent of the illumination, while the faint features in it are caused by wavefront imperfections or residual interference effects in the illumination path. The tilted fine stripes are caused by an etaloning effect in the camera window. In reference (7), the lateral position of the sample was modulated at much lower frequency than the camera frame rate of 5 kHz in order to realize a lock-in arrangement for eliminating residual wavefront imperfections that are usually present in a wide-field illumination. This allowed a direct visualization of the inherent speckle pattern caused by the sample substrate. However, as can be seen below, it is not necessary to remove the illumination inhomogeneities separately as they are also automatically eliminated in the background treatment process.

The procedure based on low-frequency sample modulation effectively reduces the frame rate. Indeed, the unit of data recorded directly on the hardware consisted of a stack of hundreds of frames in reference (7) in order to minimize the required computer storage capacity. Young *et al*. (9) improved on this approach by reducing the number of frames that were added rigidly during data acquisition to 10 and, thus, allowing for more temporal resolution in determining the contrast values (see section 2.7).

Once the rigid stacks are recorded, one can further form “batches” of many stacks during image processing to improve the SNR. To eliminate the quasi-stationary background features, two consecutive batches can either be subtracted(7; 17) or divided (9) with respect to each other, leading to a base signal about 0 or 1, respectively. In the latter case, one is subtracted from the output image to set the signal near zero once more. In both cases, the operation comes at a price of a higher pixel noise of 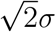 if *σ* denotes the pixel fluctuations in one batch (6; 11).

Next, to account for a finite arrival time of the particle of interest, one can perform a rolling average as a function of time, as we describe in more detail in section 2.7. We now generally refer to the combined procedure of rolling average and differential reduction of inhomogeneities as differential rolling average (DRA). Figure 2(a) shows the outcome of subtracting two batches (*B*_1_ and *B*_2_) to yield image *D*_1,2_, where the dynamic features that are new relative to the reference image can be identified (see arrows in figure 2(b)).

**Figure 2.**
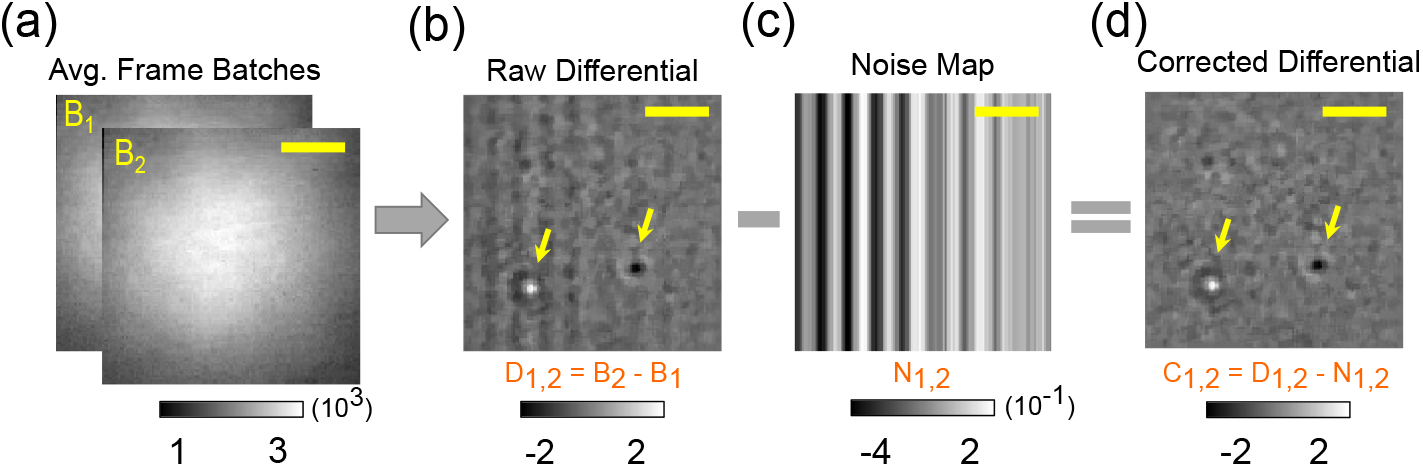
Flowchart of the fixed pattern noise (FPN) correction algorithm. Subtraction of two consecutive batches of frames *B*_1_ and *B*_2_ (a) yields a columnwise FPN (b). Two particles that are detected on the background are marked with yellow arrows. The median of each column is extracted to produce a 2D FPN (*N*_1,2_) (c), which is then subtracted from the difference map *D*_1,2_ to arrive at the corrected differential frames *C*_1,2_ (d). Scale bars correspond to 1.5 *μ*m.

### 2.4. Column-wise fixed pattern noise

As seen in figure 2(b), elimination ofthe large inhomogeneities in the image reveals weak residual vertical stripes (see also reference (7)), which we refer to as fixed pattern noise (FPN) (18). These are caused by the mismatch between the gain and the bias of the ADCs that are built in individual columns of modern CMOS imagers for improving the read-out speed. Because these parameters are not constant in time, FPN persists even after the subtraction of consecutive frames. The width and period of such a pattern dependent on the size of the FOV and frame rate. In our measurements, we typically observe a periodicity of about ten pixels, which is comparable to the size of a diffraction-limited spot (DLS), thus, complicating the identification of small particles. Previous works (9) reached a protein detection sensitivity of about 55 kDa without correcting for FPN. It is very likely, however, that FPN limits the sensitivity in iSCAT detection at some point. We now take a moment to investigate it further.

We start by assessing whether the dynamics of the observed column noise is rooted in the temporal variations of the offset or the gain of the ADC. In figure 3(a)-(c), we present the results of averaging FPNs for a bare substrate, i.e., without any nanoparticles. The bottom panels in each case display the average of pixel values in each column. The comparison of figures 3(a), 3(b) and 3c shows that averaging reduces the fluctuation amplitude. Interestingly, however, the central bright region of figure 3(c) indicates that small residual features of the illumination inhomogeneities re-appear again as a result of strong averaging.

**Figure 3.**
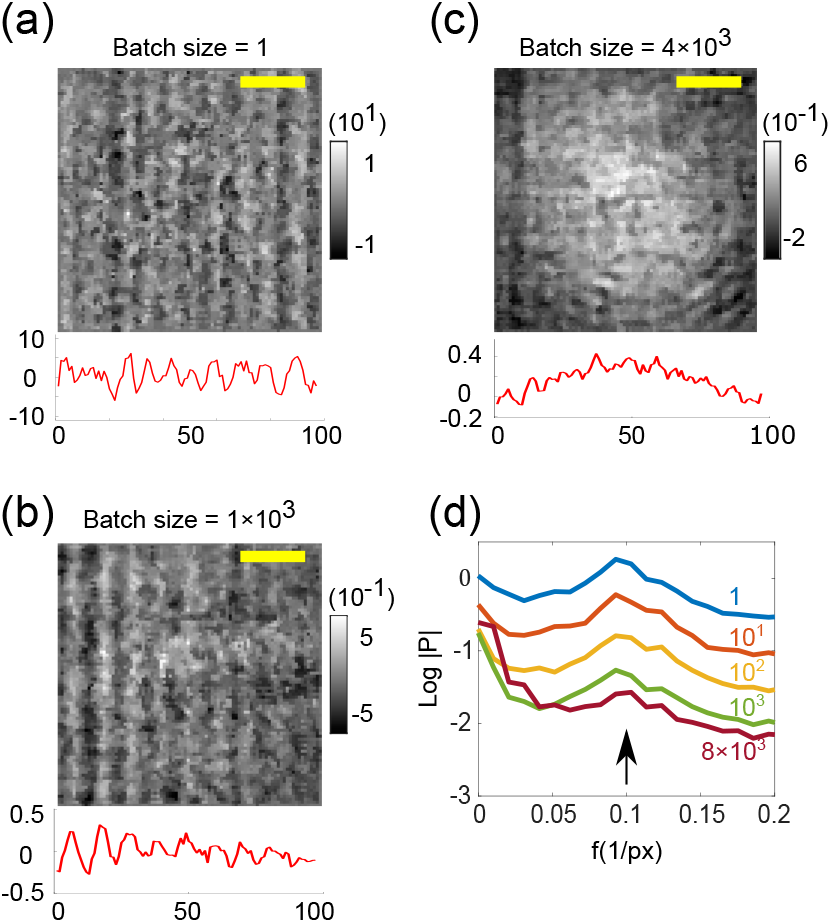
Investigating the nature of FPN. (a), (b), (c) Difference images with three different batch sizes. The pixel values in each column are averaged and and plotted underneath each image. This 1D mean signal (plotted in red) can then be Fourier transformed to obtain the periodicity of the column noise. d) The mean amplitude spectrum of bias signals for various batch sizes. The peak corresponding to the FPN periodicity is marked with a black arrow. Scale bars are equal to 1.5 *μ*m.

To quantify the FPN behavior as a function of the integration time, in figure 3(d) we plot the spatial frequency spectrum of the curves obtained by averaging the column pixels (as in the bottom panels of figure 3(a)-(c)) for a range of integration times. The main FPN periodicity is clearly represented by a peak. The comparison between the evolution of the peak height and the baseline, which is reminiscent of a 1/*f* noise, indicates that the FPN dynamics is mostly additive. In other words, it is the ADC offset that varies more than its gain. Hence, it is more appropriate to correct for FPN before the division of batches.

A variety of algorithms have accounted for FPN(19; 20; 21; 22; 23). In our work, it is important to correct for both spatial and temporal non-uniformities at the same time. Hence, we use a dedicated protocol illustrated by the flow chart in figure 2, which integrates the FPN correction into the middle of DRA. In the first part of DRA, we compute the differential frame *D*_1,2_ = *B*_2_ – *B*_1_, and then determine the median of each column. We note that we choose the median over the mean because the latter would be more susceptible to the contribution of nanoparticles that might be present in the FOV. We then generate an FPN noise map *N*_1,2_ by filling all the pixels of a given column with its median (see figure 2(c)). The noise map is further subtracted from the differential frame *D*_1,2_ to arrive at the corrected image *C*_1,2_ (see figure 2(d)). The algorithm is now applied to the remaining DRA parts.

### 2.5. Temporal behavior of the noise floor

A number of phenomena can cause temporal variations that remain in the iSCAT noise floor after DRA, thus, hindering long integration times. Examples include sample drift, vibrations in the setup, low-frequency drift of other optical components, camera pixel response, and laser beam pointing instabilities (6; 7). To investigate the temporal stability of the measurements, we monitored a 48 x 48 pixels (2.88×2.88 *μ*m^2^) region of interest (ROI) in 10^5^ consecutive frames of the corrected videos (PN and FPN) recorded from a blank sample, i.e., in the absence of any proteins. Figure 4(a) displays the standard deviations of nine pixels in the center of the FOV with 0.6 *μ*m spacing between them as a function of integration time measured at 1 kHz. While the behaviors of the individual pixels are not identical, the noise begins to level off within a few seconds for all pixels, indicating that one does not benefit from longer integration times. To obtain a global sense of the temporal fluctuations, we averaged the standard deviations of all pixels in the FOV and examine them as a function of time. The circles in figure 4(b) display the outcome of this analysis for two different frame rates of 1 kHz and 2 kHz. The comparison between the two data sets shows that fast measurements are advantageous because one can reach the required SNR before slow system instabilities dominate.

**Figure 4.**
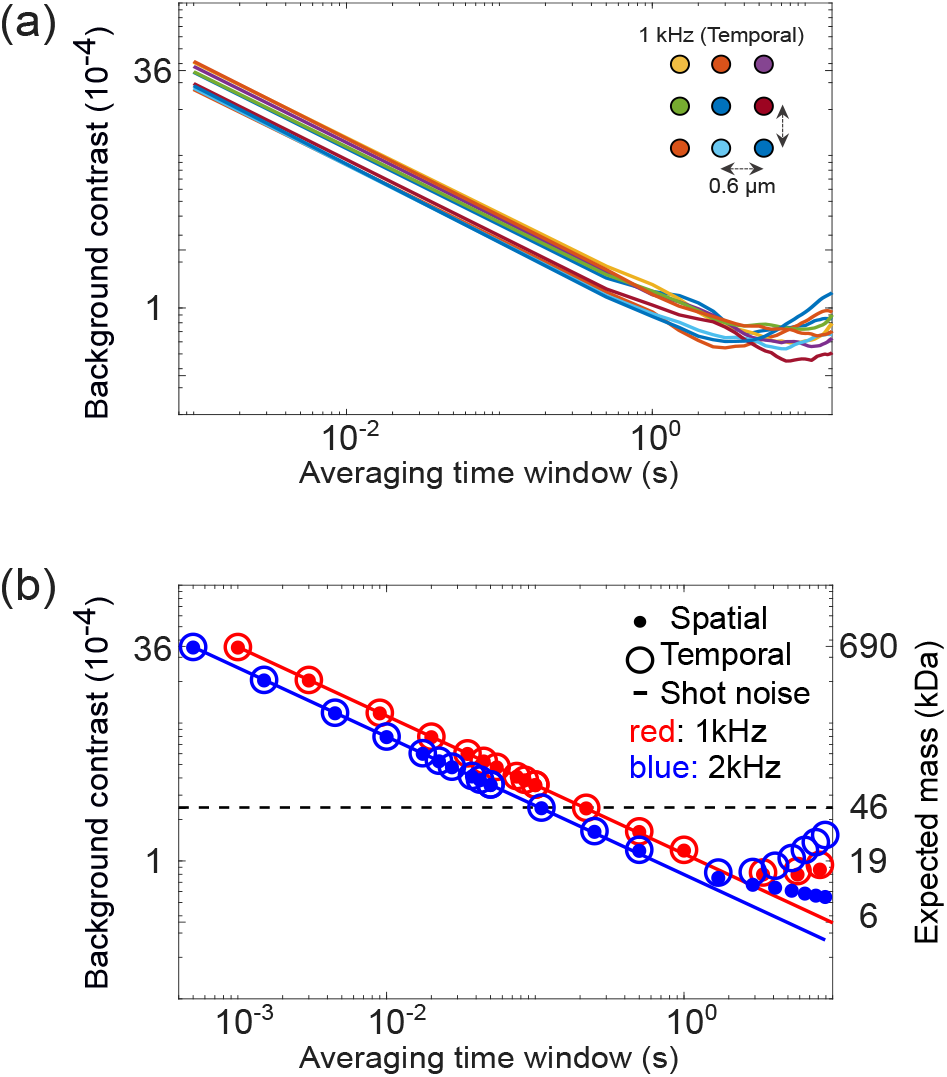
a) Standard deviation of the background fluctuations for nine pixels (neighbors are distanced by 10 pixels) as a function of batch size in DRA after PN and FPN correction. Inset shows the pixels configuration. (b) Comparison of background fluctuations for two different camera frame rates 1 (red) and 2 (blue) kHz and for temporal and spatial fluctuations. See text for details. The black dashed line depicts the theoretical resolution limit of a single 12-bit frame. The vertical axis on the right shows the expected theoretical molecular mass as a function of the averaging time window, assuming SNR=1.

A spatially fluctuating background might reduce image quality when temporal averaging is used. To characterize this aspect of a typical iSCAT image, we have also calculated the standard deviation of all the pixel values in one frame. The dots in figure 4(b) plot this quantity as a function of the integration window. We see that again averaging reduces the amplitude of fluctuations up to a time scale of the oder of 1 s, but for longer times, the noise becomes larger. We display the expected behavior of the shot noise by the solid curves and mark the single-frame digitization limit of a 12-bit ADC by the horizontal line.

The data in figure 4 suggest that after about 2 s of integration, the current experimental noise level allows one to reach an iSCAT contrast of about 1 x 10^−4^. This would imply unity SNR for the detection of single proteins with a molecular mass of 15 kDa, accessing an important class of cancer maker proteins such as cytokines (24; 25).

### 2.6. Identifying individual particles

After integration over an optimal number of frames and eliminating the unwanted background features, the remaining task is to identify individual nanoparticles and to determine their contrasts. At this point, a median filter with a size of 3x 3 pixels can be employed to remove any hot pixels that could lead to false detection. In the simplest localization method, a threshold is set to eliminate residual background fluctuations and mark features that stand out in a ROI. Here, one can take advantage of the extensive literature in fluorescence microscopy and image processing for fitting a two-dimensional (2D) Gaussian profile(26) to an experimentally measured PSF. In our case, this would provide a good approximation to the central lobe of the generally more complex iPSF(12) (see figure 1(b)). However, these common algorithms perform poorly in the presence of background inhomogeneities, especially when the signal contrast is weak. More advanced feature detection strategies have also been applied. One such example uses the Haar transform (HT) (27), which is based on dedicated square shaped wavelets and considers a locality of pixels rather than a thresholded pixel. A number of other algorithms can optimize the process for the native shape of an iPSF, which is mostly radially symmetric (28; 29). We propose and employ the Difference of Gaussian (DoG) (30; 31) for the optimal analysis of low-contrast iSCAT signals using only the main lobe of the iPSF.

In DoG, an image is first convolved with two Gaussian kernels of large and small widths and then the two resulting images are subtracted from each other. This operation acts as a bandpass filter which removes the background and extracts the sharp local maximum centered about the iPSF location (see SI. 1.2) at a radius *W*, which is 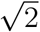 times larger than the standard deviation of the convolution. We find that DoG offers a high performance for detecting iPSFs with low SNR, localizing the center, and estimating the iPSF size, all at the same time. We note in passing that because localization algorithms are designed to detect 2D features similar to the instrument PSF, it is important to avoid spatial frequencies that correspond to the PSF width when performing pre-proccessing steps such as FPN correction (see SI. 1.3).

### 2.7. More on DRA: quantitative assessment of the contrast

At this point, potential iPSFs of the particles of interest have been identified and localized in space, reporting the number of particles and their positions. Next, the contrast of the registered iPSFs can be quantified in order to learn about the particle mass and size. In principle, one has to simply compare two consecutive batches with and without the particle to deduce its contrast. In what follows, we do this by dividing consecutive batches. However, to account for the dynamics of the particle arrival and various perturbations such as vibrations or fluidic instabilities during the integration time, it is advisable to roll batches of *k* consecutive stacks that span over the time interval between *t* and (*t* + *k*Δ*t*) (See figure 5(a)). Here, the batch size is to be set to an optimal number *k*_opt_ that is sufficient to reach a certain SNR for a given particle size of interest.

**Figure 5.**
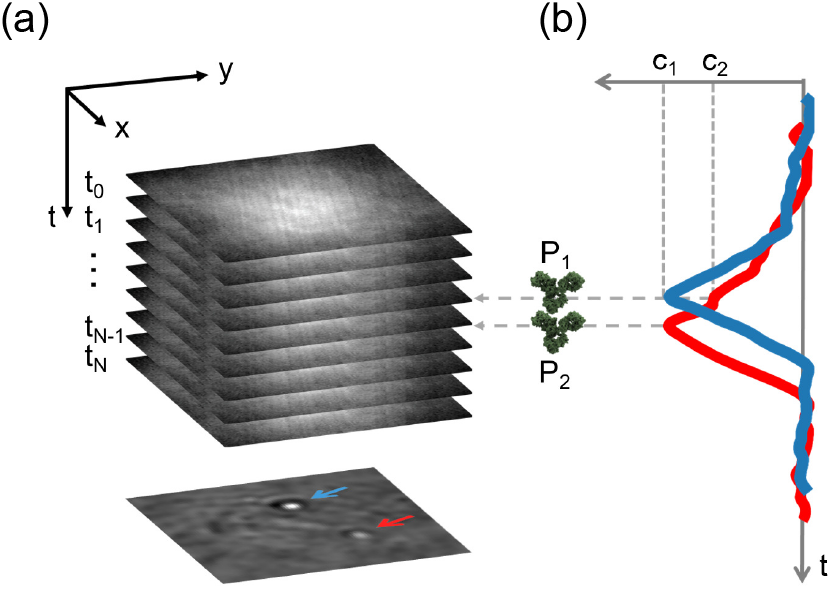
Schematic representation of the differential rolling averaging. (a) A series of batches of video frames. b) Evolution of the protein landing contrast. By shifting the consecutive batches of frames through the entire video, one reaches a V-shaped trajectory (*P*_1_ and *P*_2_) with a width equal to twice the batch size and a height that corresponds to the contrast of the particle (*C*_1_, *C*_2_). The red and blue solid lines depict the experimental contrast trajectories of two landing events for two proteins (150 kDa) indicated by arrows in (a).

It follows that as the batch division process is rolled through time, the contrast of a pixel within the particle PSF yields a V-shaped contrast profile (see figure 5(b)). We use a windowing average of 5% of the trace length to smooth the V-shaped contrast trajectories (32). Various approaches can be employed to determine the signal peak. In the ideal case, this profile has a well-defined global baseline such that its extremum is assigned to the particle contrast (9; 33). In practice, it is hard to assess the baseline of the V-profile for small SNRs, leading to uncertainties in the signal contrast. Other possibilities include reading the intersection of the two fitted lines on the sides of the V-profile or using the method of prominence (34; 35; 36). We typically monitor all three approaches.

### 2.8. Particle arrival, mobility and departure

In a realistic experiment, particles might not be present in a FOV during the whole integration time of a certain batch. As a result, the footprints of several particles in a given batch might not reach the same SNR. Furthermore, it should be kept in mind that for a camera frame rate of 1 kHz or higher, individual proteins could easily move during several frames before they bind to the surface. Indeed, a typical diffusion constant of 1-100 *μ*m^2^/s(37; 38) implies that assuming 3D diffusion, the protein may easily traverse a distance of 0.8-8 *μ*m in a typical integration time of about 100 ms.

Furthermore, considering that molecules undergo thermal agitations even after binding, a certain lateral mobility is expected. While a modest lateral jitter has no effect on the registration of large proteins, it can smear the iPSF and lower the fit quality, which has a particularly large impact on the analysis of small proteins. Figure 6 plots the contrast obtained in an analytical calculation that considers the displacement of a 2D-Gaussian PSF in discrete steps across an averaging batch. We find that small lateral motions of up to 50 nm could lead to a contrast error of about 10%.

**Figure 6.**
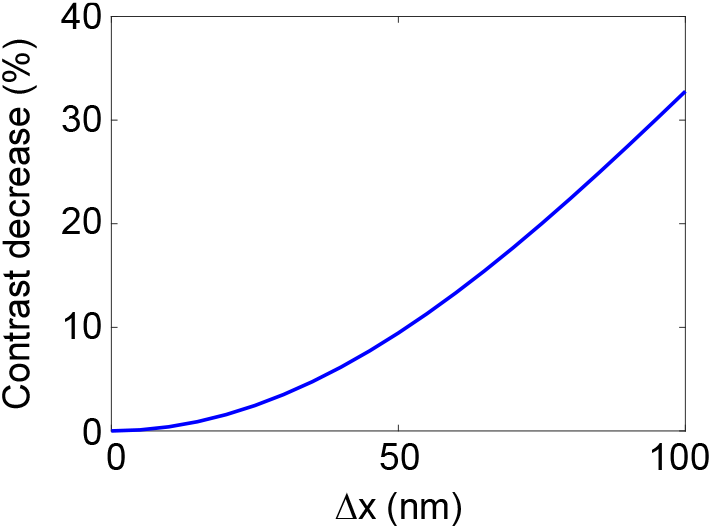
a) The influence of the particle displacement during the registration process on the measured contrast. The blue curve was generated analytically for a contrast of 1 x 10^−4^, PSF full width of 320 nm, and a displacement range of 0 to 100 nm.

Another mobility phenomenon that should be considered is the unbinding of proteins. This phenomenon can be easily identified through the reversal of contrasts that result from DRA. Figure 7 displays a map of 24 landing and 13 departure events in a FOV of 6 x 6 *μ*m^2^ on an untreated cover glass. We note that in many cases, the unbinding site is very close to a previous binding site. Furthermore, unbinding often occurs a few seconds after initial binding to the surface. The binding and unbinding rates can be controlled via the choice of surface chemistry.

**Figure 7.**
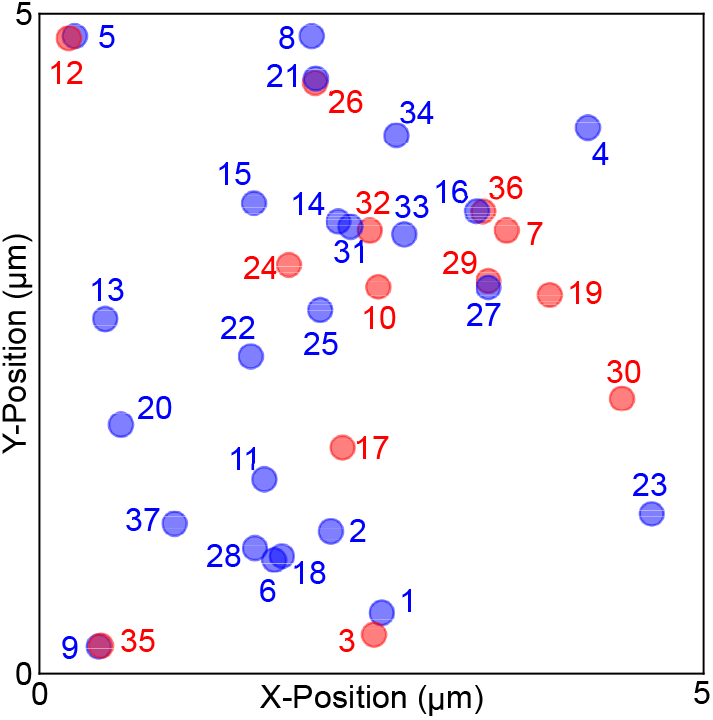
Blue and red discs illustrate the positions of binding and unbinding events for proteins of mass 66 kDa, respectively.

### 2.9. Elimination of false positive events

To ensure that we distinguish particles of interest from spurious features of a residual background, we extract the temporal behavior of each iPSF using an algorithm developed in references (39; 40). This procedure forms a full trajectory of the particle locations in a detection event if the spatial coordinates remain within a certain search range. Furthermore, the algorithm requires that a particle is not absent for longer than a certain temporal window (memory). This is especially important for iPSFs with low SNR. We choose to restrict the search range and the memory to 2 pixels (0.12 *μ*m) and 5 frames (1ms), respectively, as ideally a particle should be visible in each frame.

In addition to the above-mentioned measures, we introduce two filtering steps to ensure that we do not include unwanted signals in our detection events. First, we require that the signature of a particle is longer than a certain fraction (e.g. 60%) of the length of the DRA trace. This excludes accidental transient speckle features that might resemble an iPSF. Ideally, the length should be twice that of the averaging DRA stack size. Here, one has to keep in mind that the threshold for the trajectory length is a function of the expected SNR and of the hyperparameters setting of the localization algorithm (DoG). Indeed, high-SNR events give rise to clear full-shaped V traces, whereas those for events with a low SNR are often incomplete, especially at the opening of the V profile.

To reduce the number of false detection events, in the second filtering step we eliminate features that clearly lack radial symmetry or have a different size from the expected iPSF. Here, by using anisotropic 2D-Gaussian fitting we determine the minimum (*W*_min_) and maximum (*W*_max_) widths of the central lobe of the iPSF and require their ratio *W*_min_/*W*_max_ to be in the interval (0.7,1). We note that by imposing other requirements, e.g., on the iPSF rings (12), the selection criteria can become more restrictive.

### 2.10. Histograms of iSCAT contrast values

As in most studies, the iSCAT contrasts obtained from individual particles show statistical fluctuations. We recall that in iSCAT, the particle contrast is proportional to its polarizability, which in turn scales linearly with the particle mass if the density is assumed to be constant. Thus, the distribution of a histogram across a range of iSCAT contrasts translates to a distribution of particle mass, establishing a powerful complementary technique to mass spectrometry (9).

The histogram in figure 8(a) presents an example of this for 474 detection events of fibrinogen protein with a nominal mass of 340 kDa. We observe a prominent peak which can be fitted to a Gaussian distribution depicted by the blue curve. However, one also finds a weaker tail towards larger molecular masses. This may indicate the existence of protein aggregates for a purified sample (41). Furthermore, for a sample containing different unknown protein species (10), the measured histogram reports on the distribution of its various constituent masses. In this case, a Gaussian mixture model (GMM) can be used to identify multiple Gaussian sub-populations (42; 43). GMM is a semi-supervised probabilistic algorithm based on machine learning, which assumes the data contain a mixture of a finite number of Gaussian distributions with unknown parameters. As a statistical measure, Akaike information criterion (AIC) and Bayesian information criterion (BIC) are commonly used as two estimators to deduce the most probable number of sub-populations (42). Here, AIC is better at detecting the number of modes at the lower band limit (false negatives), whereas BIC is better at eliminating false positives at the upper band limit (over-fitting) (44; 45). We apply both AIC and BIC.

**Figure 8.**
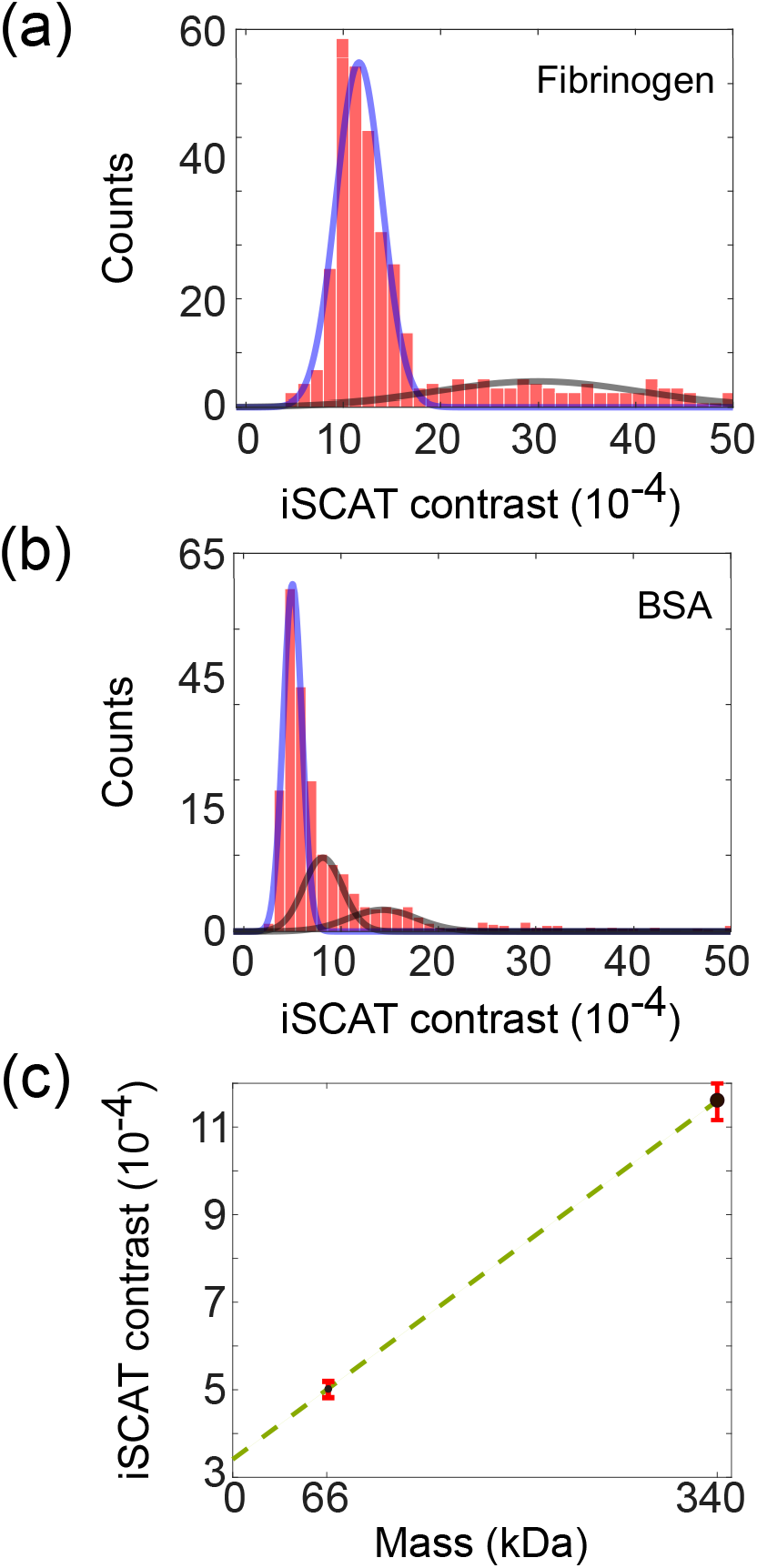
Histogram of the iSCAT contrast for binding events for samples of 340 kDa (a) and 66 kDa (b). The histogram envelopes illustrate two (a) and three (b) normal distributions obtained by GMM. The (mean±std)x10^−4^ of GMM results are 11.6±2.4and29.5±10.4for (a) and 5.0±0.90, 8.1±2.0and14.7±2.6 for (b) histograms, respectively. (c) The linear fit to the first modes of 340kDa and 66kDa histograms is depicted. The precision in determining the histogram peaks is shown by the red error bars.

Given the finite width of each Gaussian component, the ability to *resolve* the different species depends on the width of each histogram feature and the statistical robustness of the data. The spread in the contrast values may be caused by intrinsic stochastic features of the desired signal, but it may also result from sample impurities or imperfections. Moreover, systematic errors or noise in various measurement and analysis steps can influence the contrast. For example, variations of the refractive index along the optical path could lead to the modulation of the interferometric signal.

### 2.11. Precision and accuracy in mass determination

The center of a Gaussian feature can be determined with much better precision than its width just as is the case for spatial localization of single molecules in super-resolution microscopy (46). Thus, it is evident that in analogy with localization microscopy, issues such as the histogram bin size (47), the overall feature width, and the SNR play a role in the attainable precision. In the example of figure 8(a), we find *C*=(11.6±0.38)x10^−4^ for the first mode, whereas the large peak in figure 8(b) yields *C*=(5.0±0.14)x10^−4^.

Besides the issue of the measurement precision, the quantity of central importance in mass determination is the measurement accuracy, namely how close the experimental value is to the true value. Because the contrast in iSCAT depends on many parameters such as the strength of the reference field and the collection efficiency for the scattered field, it is important to establish a calibration procedure. References (7; 9) have shown that this can be obtained by measuring several well-known samples to establish a calibration ladder, much as is the case in gel electrophoresis (48). Figure 8(c) presents the general idea using the two data points extracted from figures 8(a),(b). The error bar on each data point represents the error in the fit of the Gaussian. The small intercept on the vertical axis indicates that the analysis procedure does not fully eliminate the background. Once the measurement platform has been calibrated, a new contrast measurement can be converted to a mass value. The precision in the measured contrast is then translated to an uncertainty in mass.

## 3. Summary and overview of the analysis pipeline

Figure 9 presents an overview flowchart of the signal analysis algorithm discussed in this article. Starting with a iSCAT raw video, we normalize the pixel values in each frame to the sum of all pixels in that frame to create a power normalized video. We then search for the optimum averaging batch size that, together with the FPN correction, would yield the best noise floor (See figure 4). Subsequently, particles are localized in each batch and are tracked in time. We examine the spatio-temporal characteristics of the candidate particles and filter out those which do not fulfil the selection criteria, e.g., by removing the smeared and asymmetric iPSFs. A V-shaped profile is constructed by reading the contrast of the central pixel in the PSF across the entire trajectory of a protein or by determining the amplitude of a 2D-Gaussian function fitted to its PSF. In this study, we analyzed the data with the former method. Next, GMM is used to assess the number of Gaussian populations in the data, whereby each mode represents a pure sub-population containing one type of protein or a particular oligomeric state of that protein. We can extract the mean of that mode in the contrast histogram as the representative iSCAT contrast for the protein.

**Figure 9.**
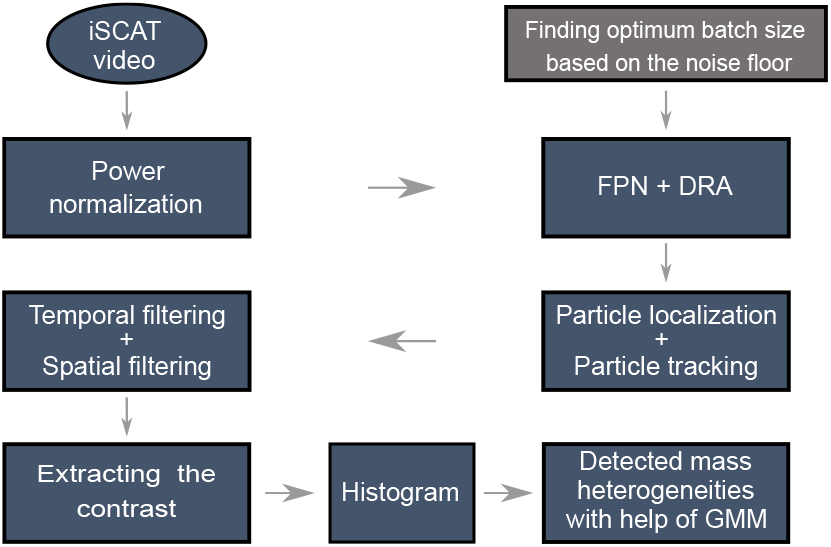
Data processing pipeline (49). See text for details.

### 3.1. Synthetic data

As we previously demonstrated, even after the sophisticated image processing procedure outlined above, there are finite background fluctuations in the image, preventing the detection of arbitrarily small proteins (see figure 10(a)). In particular, it should be kept in mind that such a background is compounded with the intrinsic shot noise of each pixel. The combination of temporal and spatial fluctuations with correlation lengths in the same order of magnitude as the microscope iPSF dictates the detection limit. To investigate this, we generated synthetic iSCAT videos and subjected them to the same analysis algorithm described in figure 9. Here, we used exemplary experimental background frames from a blank sample and superimposed PSF intensities with various contrast in a random fashion. Figure 10(b) shows the extracted contrasts as a function of the expected protein mass. The solid blue line is a fit to this synthetic library. We note that the data points extracted for cases with SNR< 3 no longer lie on the straight line because the contrast of the particle PSF cannot be reliably extracted from the noisy speckle background. Future studies will take such a synthetic analysis an important step further by considering the interference of the iPSF with the speckle background and applying machine learning algorithms to push the detection limit towards SNR ~ 1.

**Figure 10.**
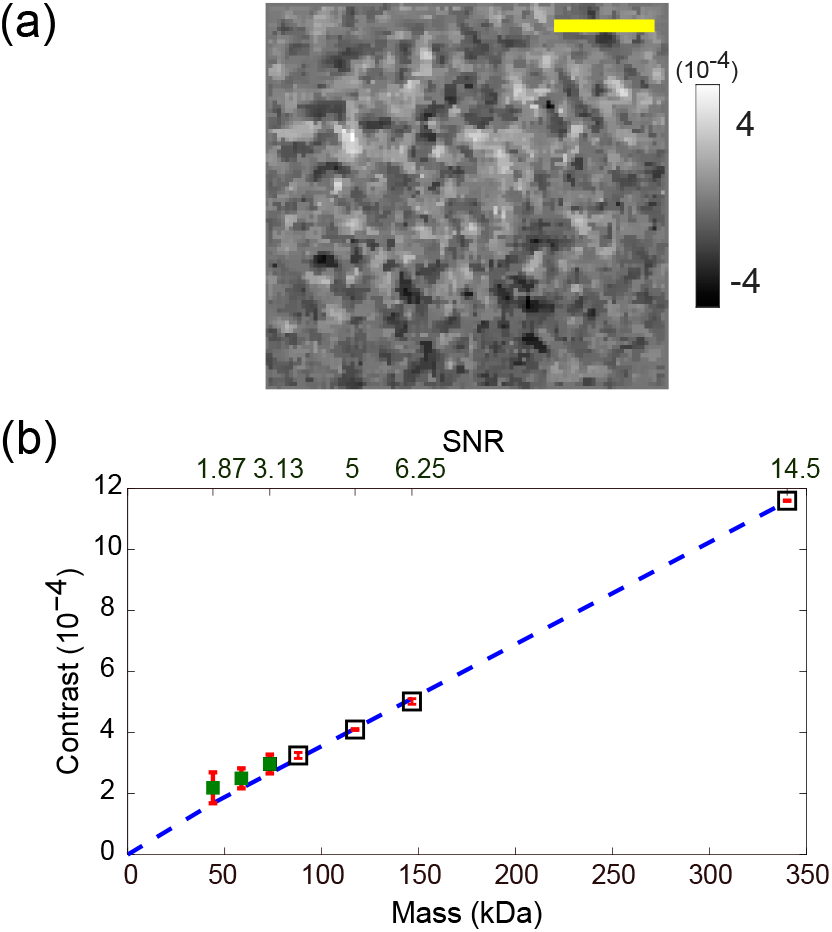
Exploring the detection limit using synthetic data. (a) An experimental example of a speckle background that remains after the whole analysis chain described in this article. Scale bar correspond to 1.5 *μ*m. (b) Contrasts of particles extracted from synthetic videos. The dashed blue line is a fit to the synthetic data excluding the three lightest particles visualized with the green squares. The error bar in each data point shows the absolute error from the fitted line by red.

## 4. Discussion and outlook

With detection sensitivity down to single proteins as small as about 60 kDa (7; 9), iSCAT has ushered optical sensing to a new horizon, well beyond the commercial gold standard based on surface plasmon resonance sensing (50). In fact, the current sensitivity is in no way limited by a fundamental issue so that there is reason to expect further advances in the near future. In this article, we have sketched the various steps involved in iSCAT protein sensing and mass determination and have identified and addressed several key factors that set the ground for future progress. Improvements can be implemented to the physical setup and measurement strategy as well as the data analysis algorithms.

On the experimental side, it will be important to examine the binding and mobility of proteins on various substrates in order to increase the detection yield and to prevent potential contrast smearing which results if a protein moves during the detection process. Employment of different camera technologies can also help improve the SNR, e.g., by providing larger well depth and better readout noise. Another interesting topic for further studies concerns a quantitative assessment of the effect of the water-glass interface on the scattering pattern of a protein and the resulting iPSF. Indeed, some recent studies have employed a spatial filter in the detection path to enhance the detected contrast (33; 51; 52). Here, it would be interesting to examine the influence of the nearfield coupling between the particle and the substrate on the angular distribution of the scattered light.

The data analysis algorithms presented in this article are provided on a python-based open-source platform (PiSCAT) (49). This analysis will be modified and extended in many different manners, especially in the wake ofa fury of machine learning techniques in image processing and microscopy (53). Furthermore, sophisticated statistical analysis of the speckle-like background and a better understanding of the spread in the contrast histograms will allow one to identify the desirable signals and eliminate false positive contributions.

## 5. Acknowledgements

We thank Alexey Shkarin for help with the camera acquisition software (pyLabLib Cam-control) and median FPN correction, Katharina König and Martin Blessing for functionalized surfaces, Tobias Utikal for the optical setup and Maksim Schwab for the mechanical components. We are also grateful to Matthias Bär for assistance with setting up the PiSCAT platform as well as Anna Kashkanova, Katharina König and Richard Taylor for valuable comments on the manuscript. This work was supported by the Max Planck Society.

## Footnotes

§ https://piscat.readthedocs.io/

## Notes

### Competing Interest Statement

The authors have declared no competing interest.

