## Supplementary material for "Optimized analysis for sensitive detection and analysis of single proteins via interferometric scattering microscopy": SI

July 2021

‡ To whom correspondence should be addressed.

### 1. Supplementary Information

#### *1.1. iSCAT contrast conversion*

##### *1.1.1. Volume to iSCAT contrast conversion*

The intensity reflection coefficient of light at the water-coverglass interface is given by:

$$R = \left(\frac{n_\mathrm{c} - n_\mathrm{m}}{n_\mathrm{c} + n_\mathrm{m}}\right)^2, \tag{1}$$

where $n_\mathrm{c}$ and $n_\mathrm{m}$ are the refractive indices of the the coverglass (1.518) and the ambient solution (1.337), respectively. Using the polarizability of the protein from equation (3) and the scattering cross section from equation (2), one can calculate the expected value of the contrast (see equation (4)) as follows:

$$C = 2\sqrt{\frac{\sigma_\mathrm{sca} \times \zeta}{DLS \times R}}, \tag{2}$$

Here, $\sigma_\mathrm{sca}$ stands for the scattering cross section, DLS denotes the area of the diffraction-limited spot of the setup and is about $\pi(100)^2\,\mathrm{nm}^2$, and $\zeta$ is the collection efficiency and is set to 35% (1). As an example, for IgG with a refractive index of 1.587 (2) and a volume of 232 $\mathrm{nm}^3$ (3), one obtains an iSCAT contrast of $7.2 \times 10^{-4}$.

##### *1.1.2. iSCAT contrast to molecular mass conversion*

Using the iSCAT contrast, one can estimate the molecular mass of the analyte. Equation (3) is used to calculate the cross-section of scattering as a function of iSCAT contrast,

$$\sigma_\mathrm{sca} = \left(\frac{C^2 \times DLS \times R}{4\zeta}\right). \tag{3}$$

Equation (4) shows the relationship between polarizability and iSCAT contrast.

$$\alpha = \sqrt{\frac{3 \times \sigma_\mathrm{sca} \times \lambda^4}{8\pi^3 (n_\mathrm{m})^4}}. \tag{4}$$

Here, it should be noted that the imaginary fraction of the polarizability is considered to be negligible compared to the real part for small proteins. Using equations (3) and (4) one can arrive at the volume ($V$) of the scatter as follows:

$$V = \frac{\alpha}{3} \times \left(\frac{n_p^2 + 2n_m^2}{n_p^2 - n_m^2}\right). \tag{5}$$

This in turn yields in the mass ($m$) of the analyte,

$$m = V\rho(M), \tag{6}$$

where $\rho(M)$ is the density of the proteins and governed by (4),

$$\rho(M) = (1.410 + 0.145 e^{\frac{-M}{13}})\ (\mathrm{g/cm^3}), \tag{7}$$

and $M$ is expressed in units of kDa. Here, we assume a constant density for the range of proteins under study. As an example, for a contrast of $2.2 \times 10^{-3}$, one obtains a molecular mass of about 400 kDa.

##### *1.2. Difference of Gaussian*

Figure 1 shows the Difference of Gaussian (DoG) (5; 6) flow chart for iPSF localization. The raw image is convolved with two 2D Gaussian kernels of widths $\sigma_1$ and $\sigma_2$, whereby the former is chosen to be comparable to the expected PSF width while the latter is set to be larger. The difference image acts as a band-edge filter and emphasizes the central part, which can be used to locate the particle. The DoG features with only contrasts above a certain threshold are localized.

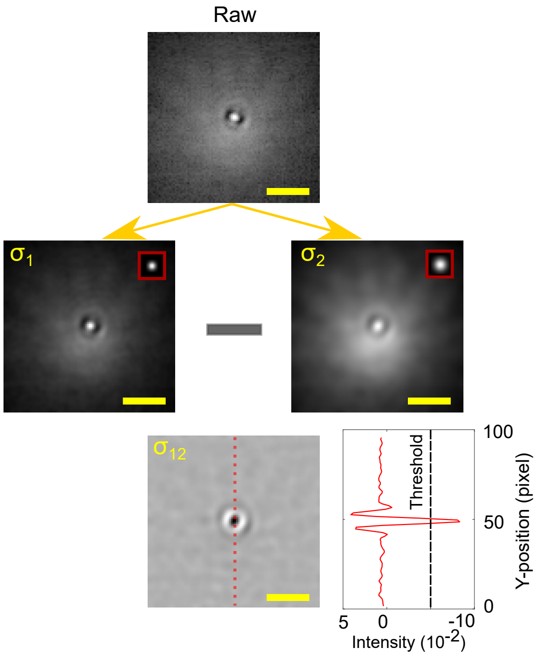


**Figure 1.** The flow diagram of DoG. Insets in the middle-row images show the Gaussian kernels for convolution in each case. Scale bars are equal to 1.5 $\mu$m.

##### *1.3. The fixed pattern noise: Effect on particle localization algorithms*

We investigate the limit to which FPN-induced spatial fluctuations in the DRA images hinder the detection of a particle. In figure 2(a), DRA images are shown in the left column before and after the FPN correction procedure. We now examine the effect of convolution with a 2D Gaussian of width equal to that of our microscope PSF, as is involved in DoG. The right column of figure 2(a) presents the outcome.

The histogram of the pixel values of the images in (a) are plotted in figure 2(b). It is evident that FPN correction helps reduce the spread in the pixel values and even to some extent the contrast offset in the convolved images. However, in case of convolved images, the reduction in the spread of the pixel values after the FPN corretion is more prominent.

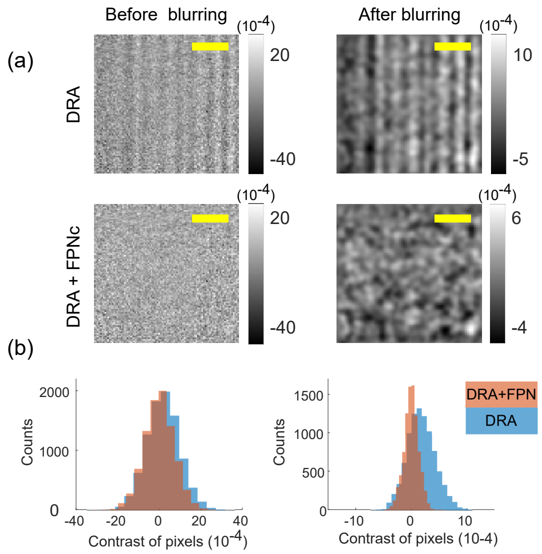


**Figure 2.** Illustration of the spatial fluctuations due to FPN pattern: a) The images in the left column are DRA images before and after the FPN correction (FPNc). Images in the right column show the results after blurring. b) The histogram of the pixel values are plotted for the DRA images before (blue) and after (orange) FPNc and overlayed in each column. For columns 1 and 2, the corresponding (means$\pm$std)$\times 10^{-4}$ are $\mu_{1,blue}$=1.8$\pm$8.0, $\mu_{1,orange}$=0.004$\pm$7.2 and $\mu_{2,blue}$=1.8$\pm$2.6, $\mu_{2,orange}$=-0.008$\pm$1.4 for each histogram. Scale bars correspond to 1.5 $\mu$m.

##### *1.4. Video Analysis: summary of the pipeline with the list of hyper-parameters*

Table 1 summarizes the list of hyper-parameters used in our analysis pipeline. In the case of the proteins with smaller molecular masses, the video recordings are done with higher frame rates and that would enable us to average larger batch sizes and improve our detection limit. The range for $\sigma$ used in the DoG algorithm was tuned based on the iPSF size while the detection threshold set for this algorithm depends on the expected iPSF contrasts.

**Table 1.** Hyper-parameters used in our localization and tracking routines: First column is the molecular mass of the protein samples injected in iSCAT experiments. Second column is the number of frames that were averaged in iSCAT videos to improve the SNR. Since recordings are done with different experimental configurations, the corresponding averaging time for each protein sample is given in the third column. The lower and upper limits of $\sigma$ values used in DoG algorithm are given in the fourth and fifth columns. This interval is sampled ten times finer. The DoG features with only contrasts above a certain threshold are localized. The threshold value is written in the sixth column.

| Mass (kDa) | Batch size | | $\sigma$ (nm) | | DoG threshold |
|---|---|---|---|---|---|
| | (Frames) | (ms) | Min. | Max. | |
| 340 | 2000 | 510 | 180 | 240 | $1.5\times10^{-4}$ |
| 66 | 2500 | 500 | 90 | 108 | $0.3\times10^{-4}$ |
